# An Engineered Live Biotherapeutic Restores Gut–Liver–Brain Axis Function by Reducing Inflammatory Pathways from Diet-Induced Stress

**DOI:** 10.64898/2026.07.10.737804

**Authors:** Andrea Verdugo-Meza, Jessica K. Josephson, Hansika Dadlani, Emad Yuzbashian, Alexander Davidson-Hunt, Ray Ishida, Sanjoy Ghosh, Deanna L. Gibson

**Affiliations:** Department of Biology, Faculty of Science, University of British Columbia, Okanagan campus; Kelowna, BC, Canada; Melius MicroBiomics Inc.; Kelowna, BC, Canada; Department of Medicine, Faculty of Medicine, University of British Columbia; Vancouver, BC, Canada

**Author notes:** Co-first authors (Andrea Verdugo-Meza, Jessica K. Josephson). **Corresponding authors:** Name: Sanjoy Ghosh and Deanna L. Gibson, Address: 3187 University Way, Kelowna, BC V1V 1V7, Canada, and.

**Keywords:** Live Biotherapeutic Products, inflammation, gut-liver-brain axis, metabolism, secondary bile acids, barrier integrity

## Abstract

Systemic inflammatory diseases can be influenced by dietary intake, with gastrointestinal dysfunction driving both metabolic and behavioural changes mirroring the altered inflammatory profile. Additionally, the use of live biotherapeutic products (LBPs) shows promise for treating metabolic and inflammatory diseases, but their efficacy is limited by poor persistence in inflamed gut environments. Designed to utilize inflammatory byproducts, the LBP EcN::*ttr* has proven efficacy in the treatment of acute and chronic colitis, however its effects on the metabolic and behavioural patterns remain uncharacterized.

We evaluated the effects of EcN::*ttr* on mice fed a proinflammatory omega-6 PUFA-rich diet. EcN::*ttr*-treated mice exhibited notable changes in the gut, including an improved expression of tight junction protein occludin, accompanied by reduced serum lipopolysaccharide (LPS) - binding protein, indicating protection against endotoxemia. EcN::*ttr* improved insulin sensitivity compared to the parental strain, associated with increased hepatic insulin receptor expression and reduced GSK3β activation and endoplasmic reticulum stress. Secondary bile acids in mice treated with EcN::*ttr* were more abundant, with increases in those associated with resolving diarrhea and bile acid detoxification. Behavioural assessment highlighted a normalization of long-term memory along with a reduction of stress management behaviours. Altogether, EcN::*ttr* restores gut–liver–brain axis function through coordinated modulation of inflammation, barrier integrity, and bile acid metabolism.

**Highlights:**

- Live Biotherapeutic Product EcN::*ttr*, designed with a fitness advantage to survive inflammation, and provides protection against a proinflammatory omega 6-rich diet
- Administration of EcN::*ttr* improved metabolic outcomes including increasing insulin sensitivity
- EcN::*ttr* increased the abundance of secondary bile acids including those that modulate bile acid detoxification
- Behavioural parameters were normalized in mice given EcN::*ttr*
- EcN::*ttr* partially normalizes gut-liver-brain axis through restoring barrier integrity, modulating inflammation and improving secondary bile acid metabolism

## Introduction

Globally, chronic inflammatory conditions are increasing in prevalence, contributing substantially to long-term healthcare burden through both primary disease and associated comorbidities. These conditions are no longer confined to aging populations, with earlier onset now frequently observed. For example, metabolic disorders such as type 2 diabetes (T2D) are increasingly diagnosed in younger individuals and are often accompanied by behavioural dysfunction, with disease severity closely associated with neurobehavioural outcomes (1). Together, this underscores the need to better understand the systemic drivers linking metabolic and inflammatory disease.

Notably, the development of T2D and other chronic diseases (like metabolic dysfunction-associated steatotic liver disease (MASLD) have been linked to diets high in the consumption of omega-6 oils, a common trait of the Western dietary pattern (WDP) that induce distinct metabolic signatures clinically (2). Seed oils typically are rich in omega-6 polyunsaturated fatty acids (PUFA) including corn-oil, have been shown to promote cardiometabolic dysfunction (3–5) and changes in behaviors in rodents (6). A potential link between the WDP and these disorders is the gut microbiome working as part of the gut-liver-brain axis, where the composition and function of the microbiome can be affected by diet. Our lab previously demonstrated that a diet rich in omega-6 PUFA is associated with dysbiosis (7) with an expansion of pathobionts known to induce inflammation (8,9). The inflammatory environment in the gut further alters the microbiome, leading to a decrease in beneficial bacteria that produce metabolites like butyrate and secondary bile acids (SBA). These metabolites are important for promoting a tolerogenic immune response, and their loss can sustain gut inflammation (10). The resulting inflammatory response can, in turn, damage the epithelium and impair its barrier function, facilitating the translocation of gut bacterial components into circulation and causing endotoxemia. Interestingly, endotoxemia has been associated with insulin resistance and the development of liver diseases (11). SBA have associated impacts on host immunity, inflammation (12) and have been linked to behavioural changes associated with anxiety, depression and memory dysfunction (13).

Recognition of the gut microbiome as a central regulator of host health has driven the development of microbiome-based therapeutics, including probiotics and live biotherapeutics (LBP), designed to promote a healthy gut microenvironment. While preclinical studies have demonstrated promise for treating intestinal disorders such as inflammatory bowel disease, translation to clinical efficacy has been poor. In particular, extra-intestinal conditions have limited studies on liver and brain-related conditions, although growing evidence links dysbiosis to extraintestinal conditions such as MASLD and anxio-depressive behaviours.

A central limitation of conventional probiotics is their inability to persist within inflamed gut environments, as they lack mechanisms to exploit inflammation-associated niches (14). This inefficiency presents an opportunity for therapeutic innovation through bioengineered therapeutics specifically designed to overcome such limitations. Inflammation generates distinct metabolic substrates, such as tetrathionate (TTR), that can be selectively utilized by bacteria equipped with appropriate metabolic pathways. Leveraging this concept, we bioengineered the probiotic *Escherichia coli* Nissle 1917 (EcN) to express the tetrathionate respiratory (*ttr*) operon, enabling persistence in inflamed environments through a competitive fitness advantage (15). This engineered strain, EcN::*ttr*, has previously been shown to reduce inflammation and promote mucosal healing in models of murine colitis (16) and porcine colitis (17). EcN::*ttr* is intended to treat populations with chronic inflammatory conditions, and as such, is developed as a live biotherapeutic product (LBP), and has been approved for pioneering human trials in Australia with the OGTR DIR license granted (license number: DIR 221).

To better represent the effects of EcN::*ttr* under WDP, we assessed the metabolic effects of EcN::*ttr* in wild-type mice fed a diet rich in omega-6 PUFA. Our findings indicate that on the WDP, EcN::*ttr* treatment but not unmodified ECN improved metabolic markers associated with insulin metabolism, endotoxemia and gut health. More importantly, EcN::*ttr* have a positive effect on anxiety and long-term memory. This novel approach demonstrates the beneficial role of LBPs in channeling the detrimental effects of diet-induced inflammation towards a positive metabolic and psychological outcome during metabolic stress via gut-liver-brain axis.

## Methods

### Ethics

Rodent experimental protocols were approved by the University of British Columbia’s Animal Care Committee per guidelines drafted by the Canadian Council on the Use of Laboratory Animals under protocols A15-0201, A19-0286, and A23-0033. All mice were assigned randomly to the experimental groups and maintained under specific pathogen-free conditions at a controlled temperature of 22±2°C, with 12h light/dark cycles, and provided with autoclaved water and food as detailed below ad libitum. Where possible, experimental staff were blinded to group assignment.

### High omega-6 PUFA diet in C57BL/6 mice

Six-week-old female C57BL/6 mice (n=9-10 per group) were purchased from Charles Rivers. Sample size was estimated from prior intraperitoneal glucose tolerance test (IPGTT) area under the curve (AUC) data using a two-sided test with α = 0.05 and 80% power. Based on the observed difference between EcN::*ttr* and the control group and pooled variance from pilot data, the required sample size was 5 animals per group. To account for potential attrition, 9-10 animals per group were used/planned.

Upon arrival, the mice were acclimated and housed until 12 weeks of age and were fed PicoLab Mouse Diet 5053. During the experiment’s duration (6 weeks), the mice were given a base diet (BioServ AIN-76A, 08822; Flemington, New Jersey, USA) supplemented with corn oil (Mazola, ACH Food Companies Inc; 60181; Oakbrook Terrace, Illinois, USA), referred to as the high omega-6 diet hereafter. This diet consisted of 80 g of the base diet and 20 g of corn oil (20% w/w). The diet was replaced every other day to limit exposure to fat oxidation. Supplemental Table 1 provides details on the composition of the omega-6 diet of which 40.8% of total energy consumed was provided by the added fat (Supplemental Table 2). Intervention with Vehicle (Luria-Bertani (LB) broth), EcN or EcN::*ttr* gavage (0.1 mL of LB broth containing 1×10^9^ to 1×10^11^ CFU/mL of bacteria, using a straight 22G, 1.5” x 1.25 mm feeding needle) occurred during experimental weeks 2 and 4. In experimental week 5, the mice were subjected to an oral glucose tolerance test (OGTT), rested for 48 hours, and then underwent an insulin tolerance test (ITT). The mice were monitored weekly, and weight records were taken once a week. Food weight was also recorded to calculate high omega-6 diet consumption during the experiment. On the last day of experimental week 6, the mice were anesthetized with isoflurane and sacrificed by cardiac puncture, followed by cervical dislocation.

### Dietary Formulation

Mice on this study were fed a 40% energy corn oil diet, high in omega-6 PUFA, to mimic the fatty acid profile common in the WDP (18). The comparison to a normal chow diet was not performed to maintain an isocaloric and isonitrogenous comparison between the treatment groups. Particularly as dietary composition has been noted to modify the microbiome independent of other factors, including body weight (19), maintaining a similar dietary profile was critical in assessing the effects of EcN and EcN::*ttr*.

### EcN and EcN::*ttr* growth conditions

*Escherichia coli* Nissle 1917 (EcN) was isolated from the commercial formulation Mutaflor (Pharma-Zentrale GmbH, 58313, Herdecke, Germany). EcN::*ttr* was engineered by cloning in a chromosomal insert of the *Salmonella enterica ttr* operon into EcN and confirmed by growth and sequencing (20). Bacteria were cultured in LB broth (per 1 L of medium: 10 g of tryptone, 5 g of yeast extract, and 10 g of NaCl, pH = 7.5) under agitation at 180 rpm at 37°C. Bacteria growth was confirmed through quantification of optical density at 600 nm (OD_600_) and plating serial dilutions on LB agar plates (1.8% agar) for colony forming units (CFU) quantification.

### Tissue collection

Blood was collected via intracardiac puncture and stored on ice until further processing (either centrifuged to collect plasma or serum). The cecum and colon were dissected and assessed for macroscopic changes documented with photographs; then, the colon was cut near the rectum, selecting the distal colon, which was cut into three sections. The most distal section was placed in 10% neutral buffered formalin and processed for histopathological analysis. The second section was stored in RNAprotect (Qiagen, 5912 PL Venlo, Netherlands) at –80°C and designated for RNA extraction and relative qPCR. The last section was flash-frozen in liquid nitrogen and used for protein extraction and analyzed via multiplex ELISA (Eve Technologies). Similarly, the liver was collected, and representative parts were collected in RNA later or flash-frozen in liquid nitrogen. Finally, the cecum was collected and immediately flash-frozen in liquid nitrogen.

### Histopathology

Fixed and processed colons were embedded in paraffin, and 5 µm sections were mounted on charged slides. The slides were stained for hematoxylin and eosin (Histo-core service by BCCHR, https://www.bcchr.ca/histologycore) and evaluated for histopathological damage by two to three observers blinded to experimental conditions, using the adapted scoring systems.

### Immunofluorescence

Colon sections were obtained from formalin-fixed tissue and embedded in paraffin. Slides were deparaffinized in xylene and then rehydrated in a gradient of ethanol. Antigen retrieval was carried out with trypsin at a concentration of 1 mg/ml at 37°C for 1 h; samples were blocked with 5% BSA-PBS for 30 min and incubated with the primary antibody (Rabbit polyclonal anti-Occludin, Cat No. GTX114949, Gene Tex; 92606; Irvine, California, USA) according to manufacturer recommendations, followed by incubation with secondary antibody (Goat anti-rabbit IgG AlexaFluor-conjugated 594-red antibody, Cat. No. A11012, Invitrogen; 88666; Waltham, Massachusetts, USA). Slides were mounted with Fluoromount containing DAPI (Invitrogen) for DNA staining of the nucleus. Images were obtained using an Olympus IX81 inverted microscope equipped with a QImaging EXi aqua Bio-Imaging camera or EVOS M5000. Images were analyzed using ImageJ. All the processing and imaging were carried out in a blind manner.

### Relative expression of genes of interest

RNA was extracted from colon and liver tissue using RNeasy Fibrous Tissue Mini Kit (Qiagen; 5912 PL Venlo, Netherlands). The cDNA was synthesized with iScript cDNA Synthesis Kit (BioRad; 94547, Hercules, California, USA). For qPCR, Sso Fast Eva Green Supermix (BioRad) and specific primers (See Supplemental Table 4) were used on the CFX manager 2.1 (BioRad). Ct values were used to calculate the relative expression of the genes of interest, using the ΔΔCt method and normalizing gene expression to reference genes (*18S* and *GADPH*) and relative abundance to the vehicle group.

### Enzyme-linked Immune assay (ELISA)

A commercial ELISA kit (Abcam ab269542 Mouse LBP SimpleStep ELISA kit) was used to quantify LPS binding protein in diluted serum (at least 1:1000 dilution). For multiplex ELISA, samples were sent to Eve Technologies for their processing for multiplex ELISA panels Mouse Metabolic Hormone 12-Plex Discovery Assay ® and Mouse High Sensitivity 18-Plex Discovery Assay ®. Direct serum samples were tested while colon samples were homogenized in RIPA buffer, and then protein content was quantified (Bio-Rad Protein Assay Kit II).

### Quantification of short-chain fatty acids (SCFA)

Gas chromatography was employed to detect acetic, butyric, and propionic acid in cecal tissues as described previously (21). A standard volatile acid mix (Sigma Aldrich; 63103, St. Louis, Missouri, USA) was used to determine the acids retention times and to construct standard curves. Data analysis was carried out in a blind manner with Chromeleon software v7.2 (Thermo Fisher Scientific; 02451, Waltham, Massachusetts, USA), obtaining the area under the peaks and here represented as the weight percentage of the total cecal tissue (g of SCFA/g of cecal tissue x 100).

### Oral Glucose Tolerance Test (OGTT) and Insulin Tolerance Test (ITT)

Mice subjected to OGTT or ITT were fasted, weighted, and then received glucose or insulin stimuli. For OGTT, mice were fasted for 6 hours and then pricked with a needle in the distal tail vein to obtain the basal glucose reading. This was followed by oral gavage of a 30% dextrose solution (2g dextrose/ kg body weight) and subsequent blood glucose readings 15, 30, 60, and 120 min post-gavage. For ITT, mice were fasted for 6h and then pricked with a needle in the distal tail vein for basal glucose reading and subsequently intraperitoneal (i.p.) injection of insulin (0.75 U of insulin/ kg body weight), reading blood glucose 15, 30, 60, and 90 min post-gavage. If glucose levels were <3.0 mmol/L, mice received an i.p. injection of 300 μL of a 20% glucose solution. Glucose readings were taken with a OneTouch Verio Flex® meter.

### Western Blots

Liver samples (20-30 mg) were processed in 500 μL of RIPA buffer (Thermo Fisher Scientific; 88666) and Halt^TM^ proteinase and phosphatase inhibitor cocktail (EDTA-Free; Thermo Fisher Scientific; 78441) and homogenized at 30 Hz for 2 minutes (Retsch Mixer Mill M400; Germany) twice with an interval of one minute in between homogenization. Then, samples were centrifuged at 15,000 × *g* for 15 minutes at 4°C, after which the supernatant was collected and stored at -20°C until further use. The RC DCTM protein assay (BioRad; 5000122) was performed using bovine serum albumin (BSA; Thermo Fisher Scientific; 23209) as a standard to measure the total protein concentration in the extracts. For SDS-PAGE and Immunoblotting, protein extracts were suspended in 1x Laemmli sample buffer (Bio-Rad; 1610747) with 50 mM of 2-mercaptoethanol (Sigma Aldrich) and incubated at 95°C for 5 minutes. Then, protein extracts (6 μg) were separated by SDS-PAGE electrophoresis (100 V for 10 minutes and 250 V for 30 minutes). The gels were then transferred onto polyvinylidene difluoride (PVDF) membranes (Bio-Rad; 1704156) by placing under 0.2 A for 30 minutes in the Trans-Blot Turbo Transfer System (Bio-Rad; 1704150). The membranes were incubated in blocking buffer (1x TBST [150 mM NaCl, 50 mM Tris-HCl, 0.1% Tween 20] with 3% BSA [Sigma Aldrich; A9418]) for 30 minutes, rinsed once in 1x TBST, and then incubated with primary antibodies for 16 hours at 4°C. Following 3x 5 min washes with 1x TBST, the membranes were incubated with secondary antibodies for 45 minutes at room temperature. The membranes were washed and incubated in Clarity Western ECL Substrate (Bio-Rad; 1705061) for 1 minute and imaged using ChemiDocTM Imaging System (Bio-Rad; 12003153). Protein bands were quantified using the Image Lab software (Bio-Rad; version 6.1.0).

### Bile Acid Quantification

Frozen cecum tissue was lyophilized, weighed and rehydrated with 3 mL of water per mg of tissue. Following homogenization, bile acids were extracted by adding 12 mL of acetonitrile/mg of tissue followed by an additional homogenization. Samples were centrifuged at 21,000 x *g* for 10 min and 20 mL of supernatant was removed and dried under a nitrogen gas stream. Dried residues were reconstituted in 100 mL of an internal standard comprised of 14 deuterium-labelled bile acids in 50% acetonitrile. High abundance bile acids were further diluted 20-fold with the internal standard. 10 mL aliquots of the sample solutions were run on an UHPLC-MS system (Agilent 1290 UHPLC and Agilent 6495B QQQ mass spectrometer (Agilent; 95051, Santa Clara, California, United States)) run in multiple-reaction monitoring mode with negative-ion detection. A Waters C18 column (2.1*150 mm, 1.7 mm) (Waters Corporation; 01757, Milford, Massachusetts, USA) was used for LC separation with a 0.01% formic acid in water mobile phase and in acetonitrile binary-solvent gradient elution as described by Han *et al.* (22). Linear regression calibration curves of individual bile acids were constructed using data from calibration solutions. Concentrations of bile acids were calculated by interpolating the calibration curves with the analyte-to-internal standard peak area ratios measured from the injected sample solution.

### Behavioural Analysis

Open field maze (OF) and novel object recognition (NOR) behavioural assessments were performed in this study using mazes from Maze Engineers (60077, Skokie, Illinois, USA). The open field maze (40cm x 40cm) was assessed for the anxiety-like behavioural profile quantified by the amount of time spent in the inner 50% of the maze (s), locomotion parameters including the total distance travelled during the test (cm), the average speed of travel (cm/s), exploratory behaviours indicated by the number of rears (supported and unsupported) during the test, and anxiety-associated behaviours including grooming episode frequency (#), total grooming episode duration (s), and number of jumps (#) during the trial. The mice were assessed for a total duration of 10 minutes whereupon they were returned to their home cage for 24 hours before the next test.

Associative memory was assessed using the Novel Object Recognition. NOR evaluated short and long-term memory retention with exposure to novel and familiar objects. Mice were habituated in the maze prior to NOR assessment and underwent a 10-minute familiarization (FAM) to acclimatize the mice to two identical familiar objects. Following FAM, mice were returned to their home cage for one hour. Mice that did not spend a minimum of 20 seconds examining the familiarization objects were disqualified from the remainder of the trial. Following the rest period, mice underwent a short-term memory assessment (STM) where a novel object was placed inside the maze in addition to the two familiar objects. Short-term memory assessment was undertaken for 10 minutes. Mice were monitored for time spent examining the novel object versus the familiar objects. Mice are then returned to their home cage for a 48-hours rest period. Following this rest period, mice re-entered the maze with one familiar object, the object present for the short-term memory assessment and a novel object. Long-term memory assessment was performed for 10 minutes and mice monitored for time spent with the novel object, the short-term memory object and familiar object.

Both short and long-term memory were assessed based on their object discrimination index. The discrimination index is calculated using the formula PI = (OoO/TT)*100 where OoO is the object of interest (Object seen twice, object seen once, novel object) is divided by the total time spent with objects during the test (TT). Short-term memory is analyzed using a Welch’s t-test and analysis of the long-term memory test is completed using a one-way ANOVA with a post hoc Tukey test to compare between the familiar object, short-term memory object and the novel object.

### Statistical analyses

GraphPad Prism software v9.0 was used for statistical analyses. All data was tested with D’Angostino and Pearson omnibus normality test (alpha = 0.05). ROUT outlier test was performed, removing definitive outliers (Q=0.1%). Comparisons between groups were made with non-parametric Kruskal-Wallis followed by Dunn’s post-test or One-way ANOVA followed by Tukey’s post-test (for multiple comparisons) for skewed and normally distributed data, respectively. Results are expressed as the mean ± SD, considering significant differences when *P*<0.05 and a trend defined as *P*<0.1. Results were plotted to display the mean ± SD for XY and bar plots, and median and interquartiles for box plots.

## Results

### EcN::*ttr* improved immunometabolic function under diet-induced stress

To determine whether EcN::*ttr* modulates systemic metabolism under dietary stress, we evaluated metabolic outcomes in mice fed an omega-6-rich diet. We have previously reported that mice fed a diet high in omega-6 PUFA, such as one supplemented with corn oil, develop metabolic disturbances like impaired glucose tolerance and insulin resistance (6). The mechanism behind these alterations appears to be linked to the proinflammatory effect of the high omega-6:omega-3 ratio in corn oil, which causes gut dysbiosis and subsequent alterations in liver, muscle, and adipose tissue (6,7). Given the importance of gut dysbiosis in the pathogenesis of metabolic dysfunction, we wondered whether a microbiome-based therapy, EcN::*ttr*, could protect mice from the detrimental effects of a high omega-6 diet. To do this, we randomized 12-week-old C57BL/6 female mice into three groups: vehicle (receiving LB broth), EcN (receiving the commercial umodified probiotic EcN in LB broth), and EcN::*ttr* (receiving the bioengineered version of EcN to persist in the inflamed gut). All mice were fed a 40% corn oil diet (high omega-6 diet) for six weeks, receiving oral gavages of the assigned treatment two and four weeks after being fed the high omega-6 diet (Figure 1A). After six weeks of being fed the high omega-6 diet, mice exhibited similar weight changes (Figure 1B) and responses to the oral glucose tolerance test (Figure 1C). However, the insulin tolerance test revealed that mice treated with EcN::*ttr* had greater insulin sensitivity when compared to EcN-treated mice, as glucose levels dropped almost 50% from basal levels at 15 minutes post-insulin injection (Figure 1D, left). This was reflected in a greater area over the curve (AOC) using the baseline glucose levels (p=0.0006, Figure 1D, right).

**Figure 1:**
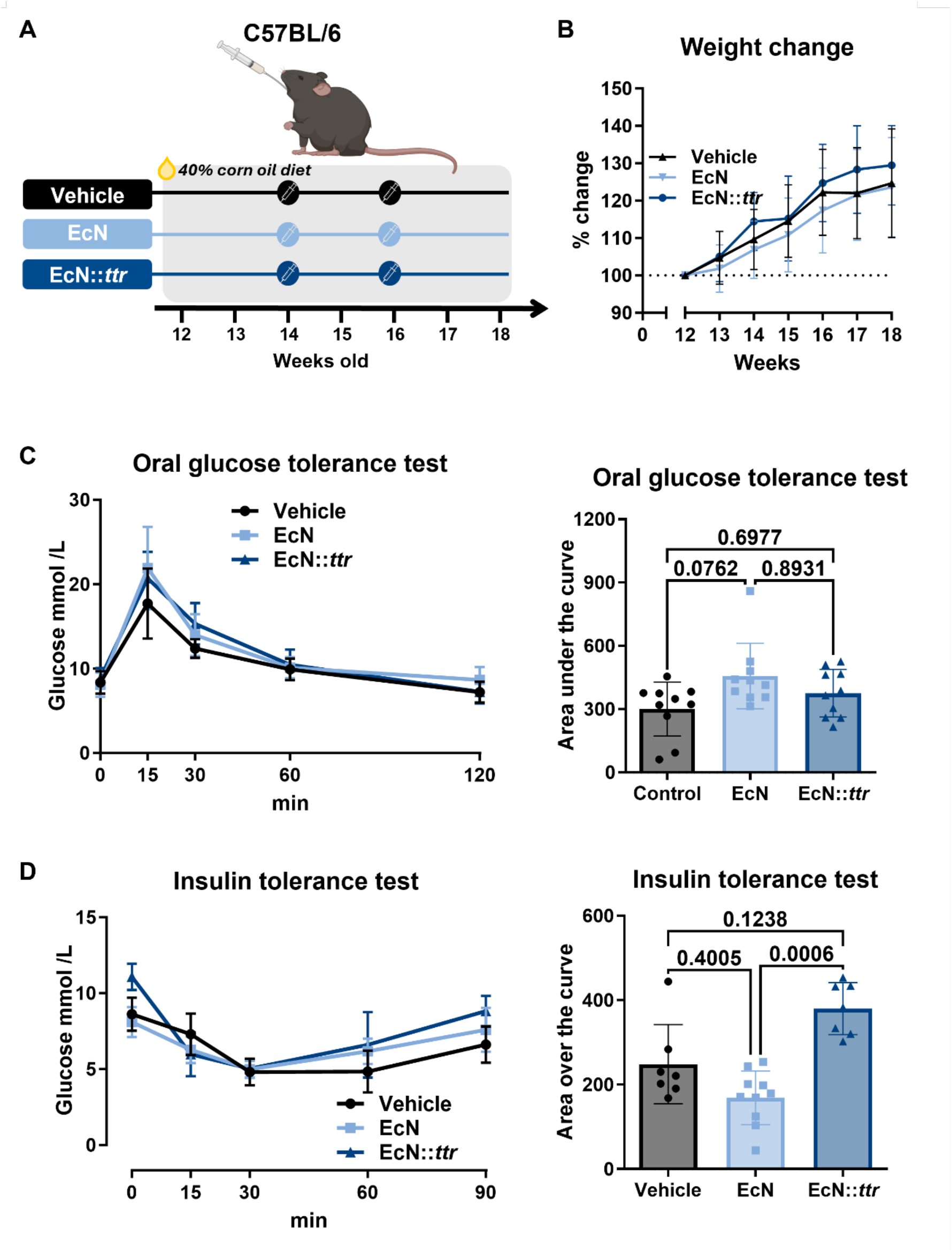
Mice fed a high omega 6-diet and treated with EcN::*ttr* exhibit greater insulin sensitivity in mice fed a high omega-6 diet. (A), experimental approach; (B), weight change; (C), oral glucose tolerance test showing glucose clearance over time (left) and area under the curve (right); (D), insulin tolerance test showing glucose levels over time (left) and the area over the curve for each group (right). n = 8-10 mice per group. Data was analyzed with Kruskal-Wallis. Adjusted *P*-values are shown in the graph.

To understand what was driving the enhanced insulin sensitivity in EcN::*ttr-*treated mice, we examined liver metabolic markers. Liver metabolism plays a key role in metabolic health and is closely linked to the gut via the gut-liver axis. We focused on the insulin pathway, as we hypothesized that EcN::*ttr* could improve the gut-derived metabolites that exhibit extra-intestinal effects. Therefore, we assessed the protein expression of the insulin receptor and downstream intracellular targets in liver protein extracts (Figure 2A). We found a modest trend towards increased insulin receptor expression (p=0.0691 EcN::*ttr* vs Vehicle, and p=0.0705 EcN::*ttr* vs EcN, Figure 1B, left), which may be consistent with enhanced glucose uptake through insulin-dependent pathways. Although there was no increase in the activation of the downstream target Akt (Figure 2B, center), we noted a modest decrease in the expression of Akt’s downstream target pGSK3β (p=0.0671 compared to vehicle, Figure 2B, right). Regulating glucose homeostasis, inflammation and ER stress, pGSK3β is a target of therapies such as Tideglusib, providing both a neuroprotective and anti-inflammatory effect. Additionally, we evaluated liver markers related to fatty acid metabolism as the 40% corn oil diet increases hepatic fat metabolism. Although we observed a rising trend for increased fatty acid uptake receptor CD36 and downstream transcription factors PGC-1, PPARα, and PPARγ these changes did not reach statistical significance (Figure 2C). Similarly, we observed modest decreases in gene expression of SREBP1c, involved in cholesterol synthesis. However, we did see changes in endoplasmic reticulum (ER) stress markers uXbp1 and sXbp1, noting a significant decrease in the sXbp1/uXbp1 ratio, which has been linked to decreased ER stress in the liver, as the spliced form of Xbp1 (sXbp1) increases with metabolic stress related to protein unfolding (p=0.0473 compared to vehicle mice, Figure 2D). This suggests that EcN*::ttr*-treated mice are more protected from liver ER stress, with improved insulin signalling under a high omega-6 PUFA diet. Together, these findings demonstrate that EcN::*ttr* enhances metabolic function in the context of diet-induced metabolic dysfunction.

**Figure 2:**
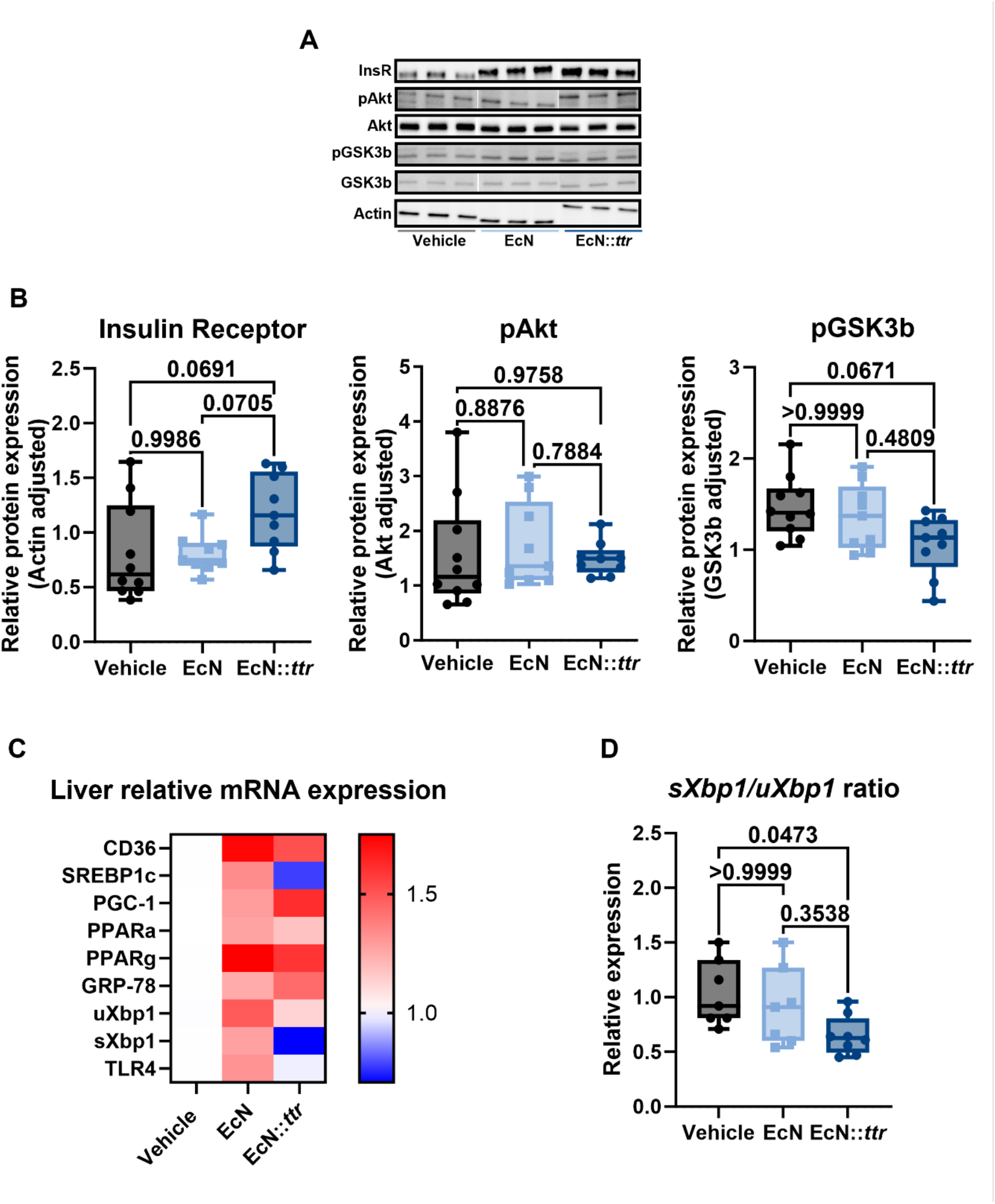
EcN::*ttr* has an impact on liver metabolic markers in mice fed a high omega-6 diet. (A), representative immunoblots of liver proteins; (B), liver protein expression of insulin receptor (left), pAkt (center) and pGSK3β (right); (C), liver relative mRNA of diverse markers associated with lipid metabolism, endoplasmic reticulum stress and LPS receptor; (D), Liver relative expression ratio of sXbp1 and uXbp1. n = 7-10 mice per group. Data were analyzed using Kruskal-Wallis. Adjusted *P*-values are shown in the graph.

### EcN::*ttr* restores gut barrier integrity and reduced endotoxemia associated with local and systemic inflammation

Given the central role of the intestinal barrier in regulating systemic inflammation, we next assessed gut epithelial integrity. We quantified the levels of LPS-binding protein in serum to further explore whether EcN:*ttr* treatment was preventing endotoxemia. Corn-oil fed mice treated with EcN::*ttr* had modest decreased levels when compared to vehicle-treated corn-oil fed mice (p=0.0736 compared to vehicle, Figure 3A). Next, we evaluated the macroscopic and histological changes in the colon (Figure 3B). We did not find macroscopic changes in the colon of mice across the different groups. Assessing histological scores (Figure 3C) revealed EcN::*ttr* was able to preserve crypt architecture while maintaining greater crypt length compared to both the vehicle and EcN groups (p=0.0146 and p<0.0001 respectively). More importantly, we found that mice receiving EcN::*ttr* had higher levels of butyric acid in the cecal contents compared to those mice receiving the vehicle treatment (p=0.0391, Figure 3D). Butyric acid, a short-chain fatty acid associated with modulation of epithelial barrier integrity and regulation of inflammation in the GI tract. We observed a notable impact on colonic inflammation in mice that were treated with EcN::*ttr* (Figure 3E) with lower levels of the pro-inflammatory cytokines IL-1β (p=0.0360 compared to EcN), IL-12 (p=0.0706 compared to vehicle), and IL-17A (0.0579 compared to EcN) when compared to the other treatment groups. EcN::*ttr*-treated mice displayed a decrease in other notable cytokines including IL-5 (p=0.0215 compared to EcN), and KC (p=0.0138 compared to EcN), involved in T cell activation and neutrophil recruitment, respectively.

**Figure 3:**
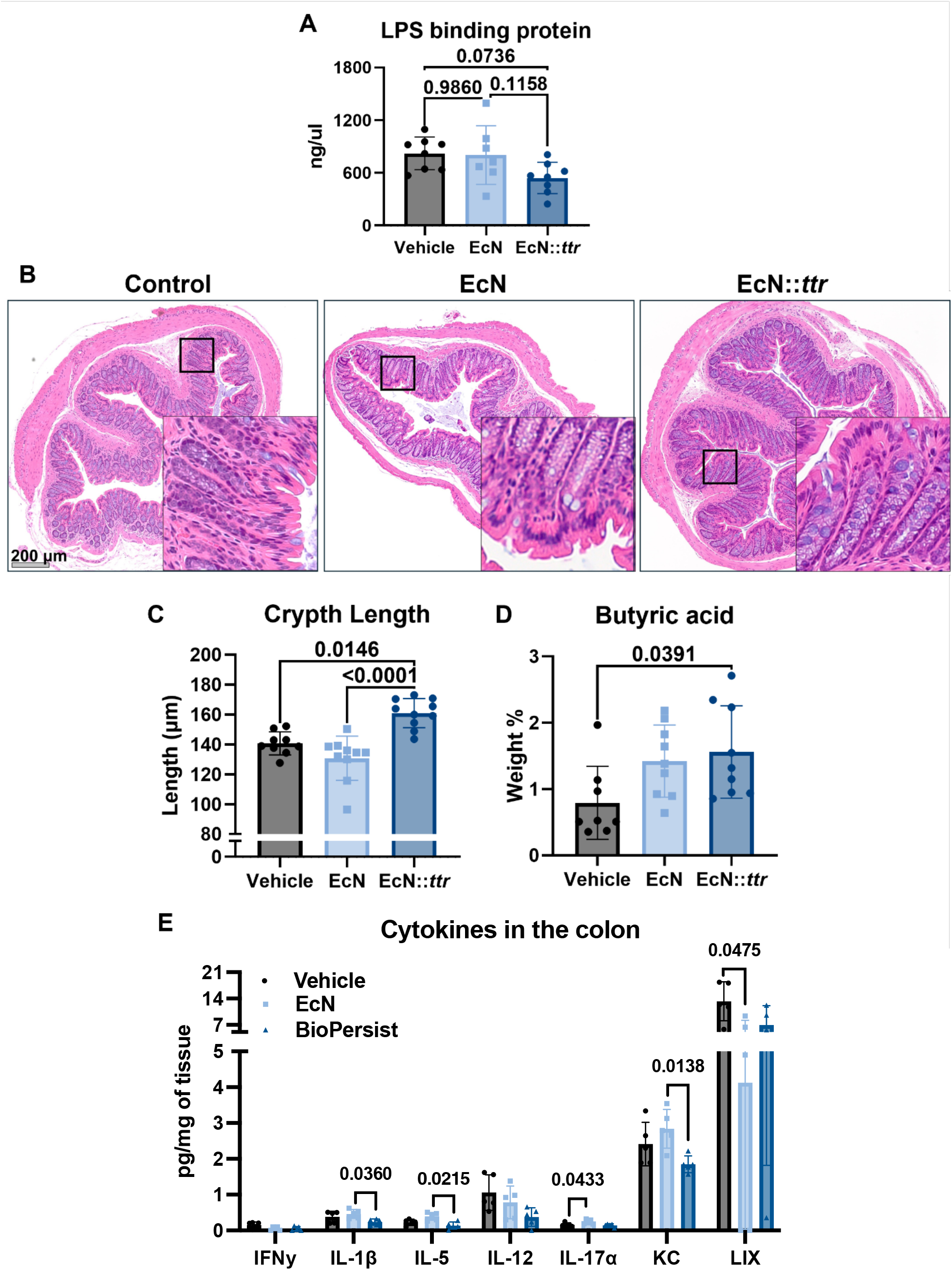
EcN::*ttr* improved serum and colon markers in mice fed a high omega-6 PUFA diet. (A), serum levels of LPS-binding protein; (B) representative cross-section of the colon; (C), crypt length among; (D), butyric acid levels in the cecum; (E), cytokine levels in colon tissue. n = 5-10 mice per group. Data were analyzed using one-way ANOVA (D) or Kruskal-Wallis (A, C, E). Adjusted P-values are shown in the graph.

Finally, to evaluate the effects of EcN::*ttr* on the epithelial barrier and explain the reduced levels of LPS-binding protein in serum, we assessed the tight junction barrier protein occludin (Figure 4). EcN::*ttr* treatment resulted in improved localization and expression patterns of the tight junction protein consistent with enhanced barrier function (23). These findings suggest that EcN::*ttr* mitigates diet-induced barrier dysfunction and limits systemic exposure to microbial-derived inflammatory signals.

**Figure 4:**
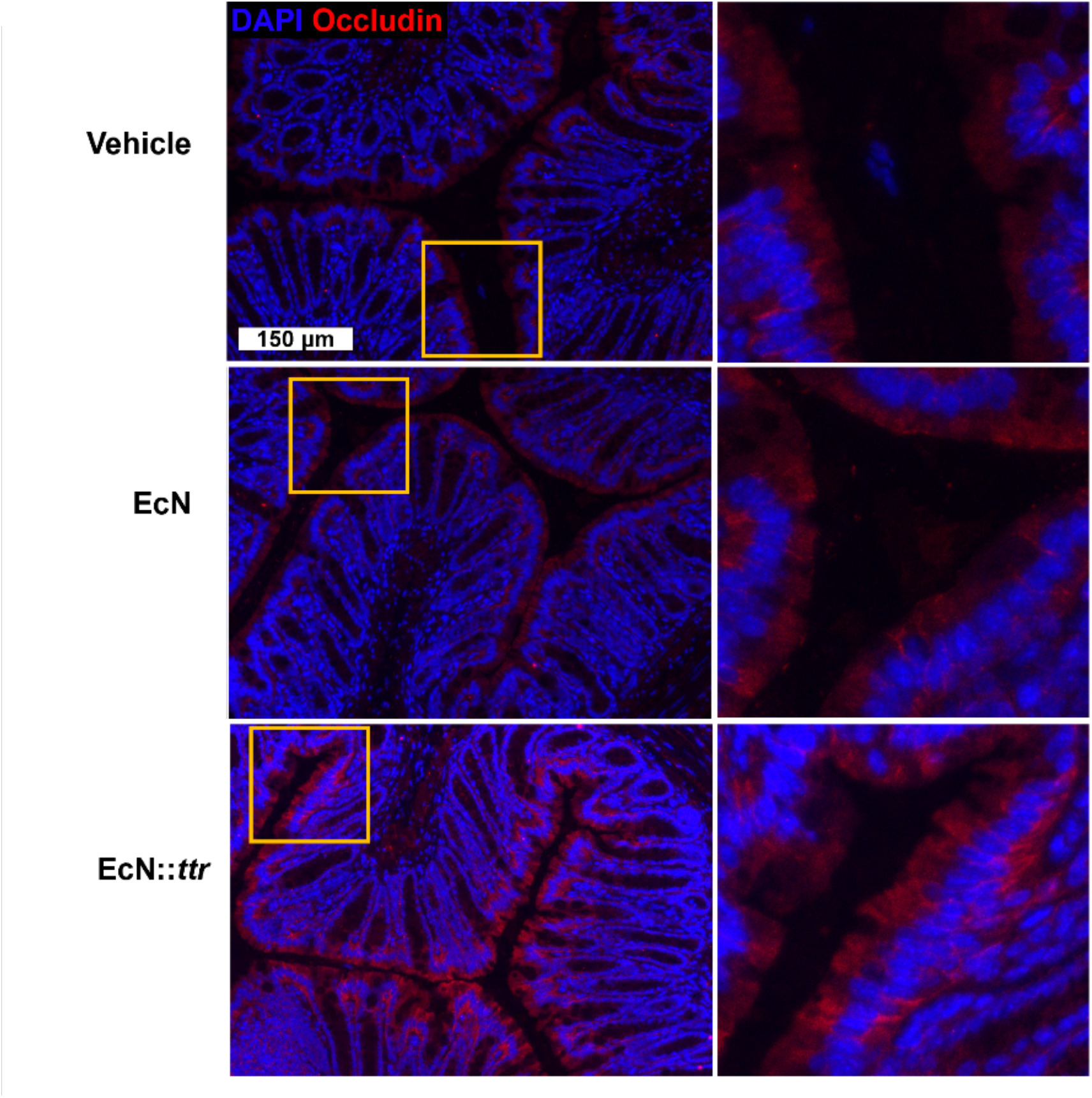
EcN::*ttr* increases the organizational expression of the barrier protein occludin in mice fed a high omega-6 PUFA diet. Representative immunofluorescence images of formalin-fixed paraffin-embedded cross-sections of the distal colon from Vehicle-, EcN-, and EcN::*ttr*-treated mice. Fluorescence tagged DAPI (blue) and occludin (red) tissues with magnified sections highlighted (right) showing improved expression of the barrier protein occludin in the epithelial layer for EcN::*ttr*-treated mice with an increase in fluorescence signal.

### EcN::*ttr* reshapes bile acid composition towards a beneficial profile

Previous untargeted metabolomic assessment of EcN::*ttr* have indicated alterations in the bile acid network as a method of action (24). To investigate whether the observed microbial and barrier changes were associated with altered metabolic signaling, we performed bile acid profiling from the cecal contents. Bile acids, especially hydrophobic bile acids, can damage the mucosal barrier and be toxic to epithelial cells (25). Therefore, the host has mechanisms to detoxify bile acids through sulfation and glucuronidation (26). EcN and EcN::*ttr* treatment decreased primary bile acids chenodeoxycholic acids compared to vehicle mice (p=0.0240 for EcN and p=0.0169 for EcN::*ttr*) (Figure 5A) and β-muricholic acid (p=0.0536 EcN::*ttr* compared to vehicle), which could indicate a greater microbial activity in transforming primary bile acids into secondary bile acids.

**Figure 5:**
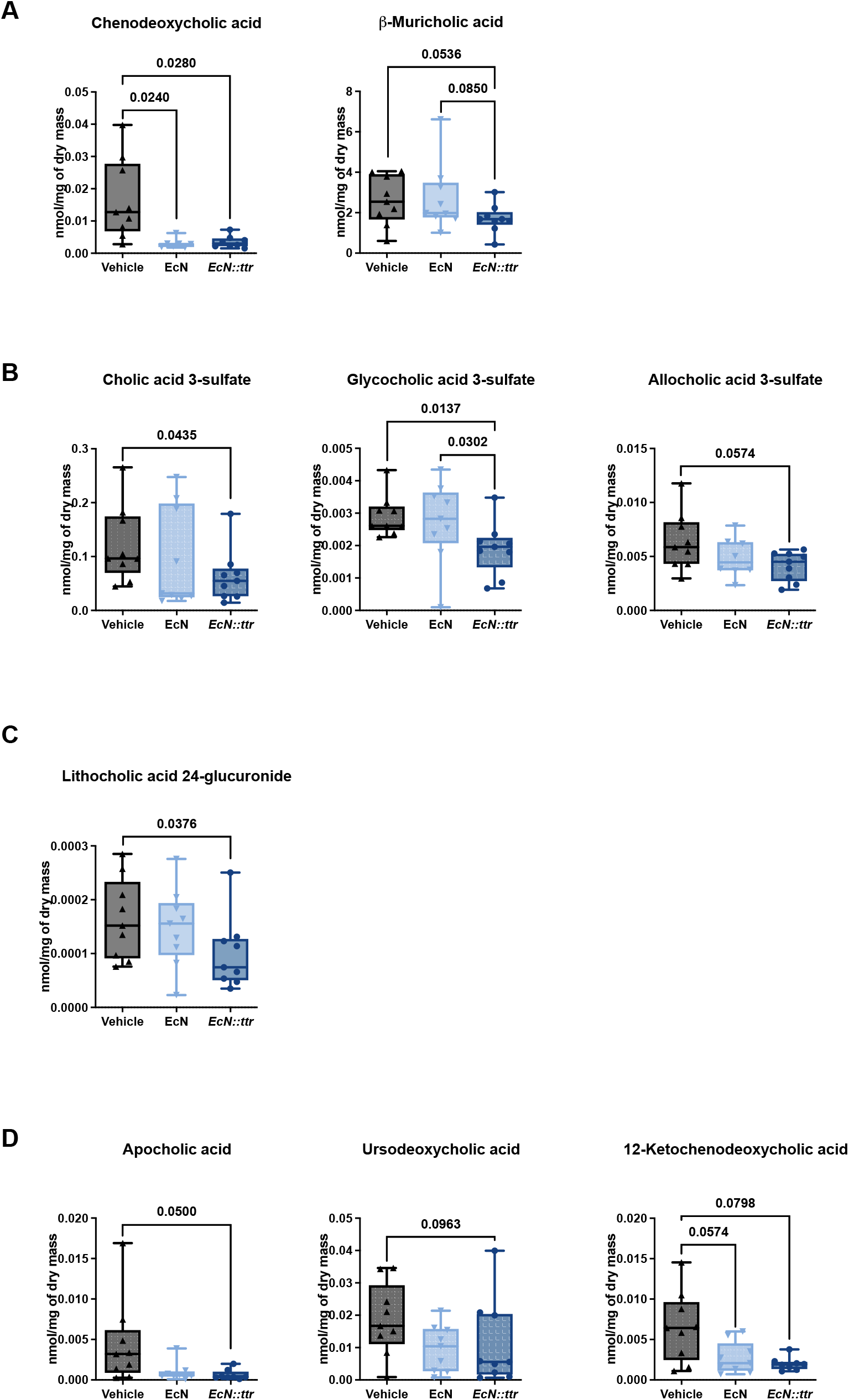
EcN::*ttr* alters the bile acid concentrations in mice fed a high omega-6 PUFA diet. Bile acid concentrations (nmol/mg of dry mass) isolated from cecal contents (n=9 per group) with changes highlighted in (A) primary bile acids, (B) sulfated bile acids, (C) glucuronidated bile acid, and (D) diarrhoea-associated bile acids. Data was analysed using Kruskal-Wallis with post-hoc Dunn’s multiple comparisons test. Adjusted *P*-values are shown in the graph.

EcN::*ttr*- treated mice showed decreased levels of sulfated bile acids compared to the vehicle treated mice (cholic acid 3-sulfate (p=0.0435), glycocholic acid 3-sulfate (p=0.0137) and allocholic acid 3-sulfate (p=0.0574)) (Figure 5B), a glucuronidated bile acid (Figure 5C) (lithocholic acid 24-glucuronide (p=0.0372 EcN::*ttr* to vehicle), and diarrhea-associated bile acids (Figure 5D) (apocholic acid (p=0.0500 EcN::*ttr* compared to vehicle), 12-ketochenodeoxycholic acid (vehicle compared to EcN (p=0.0574) and EcN::*ttr* (p=0.0798) and ursodeoxycholic acid (p=0.0963 EcN::*ttr* compared to vehicle)), which could indicate a more balanced, detoxified bile acid pool. Although EcN-treated mice also presented some significant changes in bile acids including hyocholic acid and deoxycholic acid 3-sulfate (See Supplemental Figure 1) the diversity pool is smaller.

In support of previous results, our lab had reported that in the DSS model of colitis, mice treated with EcN::*ttr* were more efficient at recycling primary bile acids, as indicated by a reduced pool of primary bile acids in the cecum (14). Moreover, here we found that EcN::*ttr* differentially decreased bile acids linked to mucosal permeability, including the chenodeoxycholic acid, deoxycholic acid, and the bile acid precursor 7α-hydroxy-3-oxo-4-cholestenoic acid. Regarding bile acids with metabolic effects, EcN-treated mice showed decreased levels of hyocholic acid, which has been suggested as an early marker of metabolic dysfunction (pre-diabetes) (27). Conversely, EcN::*ttr*-treated mice exhibited decreased levels of β-muricholic acid, a marker of suppressed hepatic lipid accumulation considered beneficial in rodent models of MASLD.

### EcN::*ttr* normalizes long-term memory and reduces self-soothing in an anxious environment

We next examined whether systemic and intestinal improvements were associated with changes in behaviour. Within this study, only the long-term memory of the mice treated with EcN::*ttr* expressed a typical focus on the novel object (Figure 6A EcN*::ttr* mice showed significantly increased preference index to the novel object compared to the object seen once (p=0.0007) and the object seen twice (p<0.0001)), whereas both the vehicle and EcN treated mice showed an abnormal preference for objects previously seen. This suggests that EcN*::ttr* administration may be a possible avenue for addressing behavioural pathologies present under systemic inflammation. When assessing anxiety-like behaviours, the OF maze showed a significant decrease in self-soothing in the EcN::*ttr* group alone through grooming time (Figure 6B; p=0.0192 compared to vehicle and p=0.0580 compared to EcN) and the number of grooming sessions (Figure 6C; p=0.0074 compared to vehicle and p=0.0123 compared to EcN). Attributed to a self-management to reduce anxiety (28), the reduction in grooming can be associated with a reduction in overall anxiety felt by the mice in the EcN::*ttr* group. Overall, our findings suggest that EcN::*ttr* offers a protective effect in the gut of mice when fed a high omega-6 diet, as inflammation and related damage decrease, resulting in subtle extraintestinal changes that remove cellular stressors and lead to an improved insulin response. Additionally, the impact of these subtle changes results in a shift in the gut-liver-brain axis resulting in normalized behaviours in both memory and anxiety-like behaviours.

**Figure 6:**
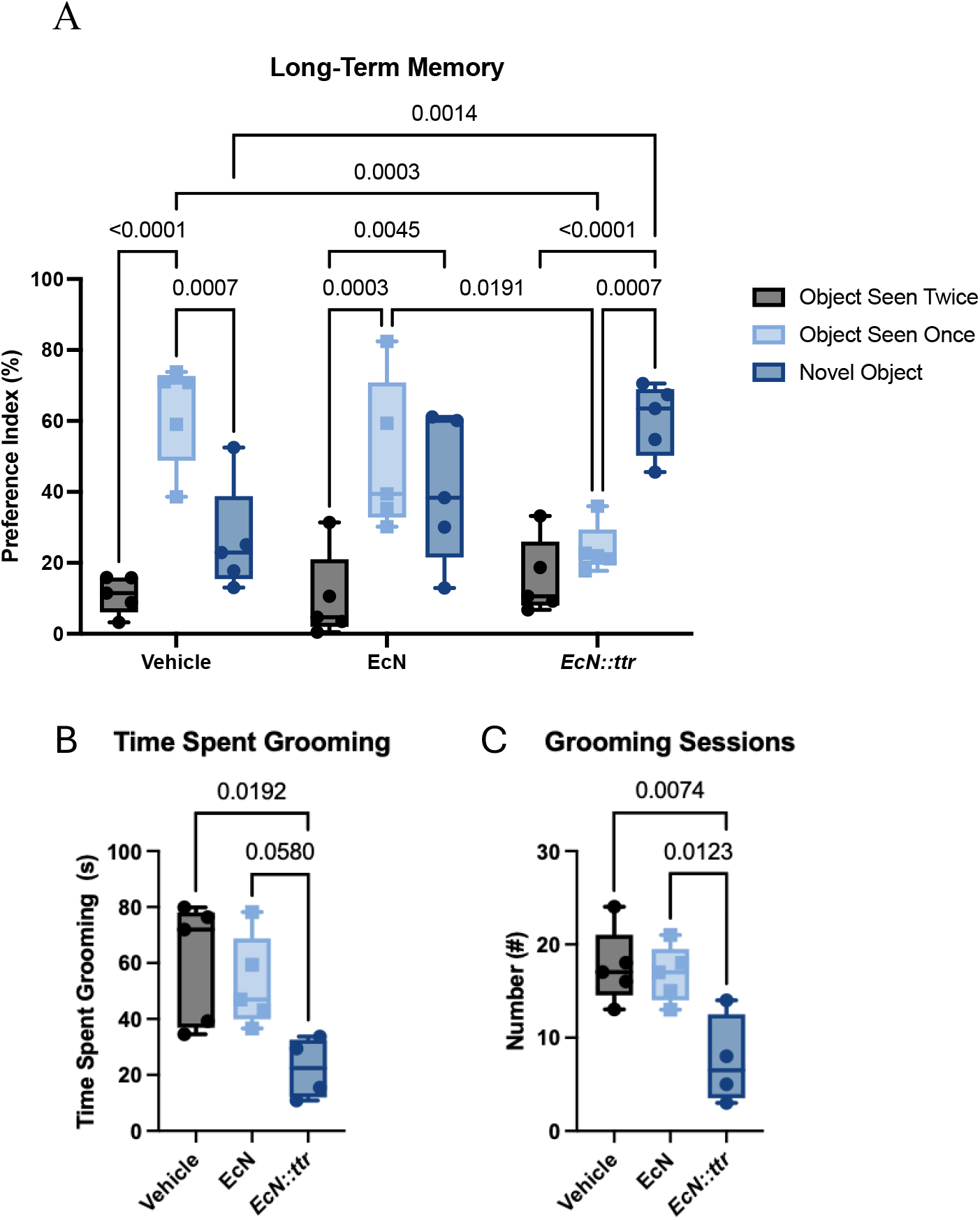
EcN::*ttr* normalizes long-term memory and reduces self-soothing behaviour in mice fed a high omega-6 PUFA diet. Vehicle- (n=5), EcN- (n=5), and EcN::ttr- (n=4) treated mice were assessed for (A) long-term memory using the novel object recognition maze and (B-C) anxiety-like behaviours using the open field maze. Normalized long-term memory was displayed for EcN:*ttr*- group only whereas EcN::*ttr* grooming behaviours showed significant decreases in B) time spent grooming and C) the number of grooming episodes.Video was analyzed by EthoVision 17. Data was analysed using a one-way ANOVA with post-hoc Tukey test.

## Discussion

Dysregulated intestinal inflammation has been associated with chronic inflammatory conditions that can lead to metabolic and mental disorders. The resulting inflammation-induced dysbiosis displaces beneficial microbes, leading to an increase in opportunistic pathobionts that can trigger inflammation. Therefore, therapies aimed at restoring the gut microbiome’s symbiosis with the host could protect not only against intestinal inflammation but also from metabolic and mental diseases. This study demonstrates that a bioengineered live biotherapeutic, EcN::*ttr*, restores multiple facets of host physiology disrupted by a pro-inflammatory diet, including metabolic function, intestinal barrier integrity, bile acid composition, and can alter behavioural patterns in a similar manner to pharmaceutical interventions. These coordinated effects support a model in which targeted modulation of the gut microbiome can influence systemic physiology through the gut–liver–brain axis.

A central finding of this work is that EcN::*ttr* improves insulin sensitivity under diet-induced stress, accompanied by increased hepatic insulin receptor expression and reduced activation of GSK3β and endoplasmic reticulum stress pathways. The consumption of a diet rich in omega-6 and low in omega-3 PUFA has been linked to intestinal inflammation in mice, (29,30) and as a risk factor for developing ulcerative colitis in humans (31). The increased omega-6:omega-3 ratio can skew the production of prostaglandin-2 and leukotriene-4, which have pro-inflammatory properties (32). This could lead to an increase in pro-inflammatory cytokines, such as IL-1β, IL-6, and TNF-α, which can suppress insulin signalling (33). EcN::*ttr* decreased the expression of pro-inflammatory cytokines in the colon, and this correlated with increased insulin sensitivity. Similarly, in the liver, we observed a subtle increase in the expression of the insulin receptor and a decrease in the activation of GSK3β. This is relevant, as GSK3β inhibitors are used for the treatment of neurodegenerative diseases have potential therapeutic effects in colitis (34). Another intracellular aspect that EcN::*ttr* modulated was Xbp1, an unfolded protein response mechanism that serves as a marker of ER stress. EcN*::ttr* decreased sXbp1/uXbp1 ratio in the liver, indicating decreased activation of the unfolded protein response in relation to the spliced version of Xbp1 (sXbp1) compared to the unspliced version (uXbp1). This suggests that the targeted gut microbiome intervention prevented the unfolded protein response triggered by ER stress in the liver by reducing gut-derived inflammatory signals. EcN::*ttr* promoted higher levels of cecal butyrate, a known anti-inflammatory bacterial metabolite (30). These results suggest that microbiome-targeted interventions can influence core metabolic signaling pathways beyond the intestine. While metabolic dysfunction is often treated as a host-intrinsic process, our findings reinforce the concept that microbial and intestinal factors are key upstream regulators of systemic metabolism.

We further show that EcN::*ttr* enhances intestinal barrier integrity, as evidenced by improved occludin’s pattern of expression and reduced circulating LPS-binding protein. This indicates protection against endotoxemia, a known contributor to insulin resistance and liver pathology (35). Restoration of barrier function likely represents a critical mechanism linking microbial modulation to systemic metabolic improvements, as reduced translocation of microbial products can dampen chronic low-grade inflammation. Indeed, rodent studies have demonstrated that a high-fat diet increases intestinal permeability, with alterations in tight junction proteins such as ZO-1 (36), and that injection of LPS can recapitulate endotoxemia and the metabolic disturbances associated with a high-fat diet (11). Although we found some protection, previous work on EcN::*ttr* in aged mice fed the same high omega-6 diet showed more significant findings, such as protection against weight gain, improved glucose clearance after an intraperitoneal glucose tolerance test, and increased plasma insulin levels (37). In our study, mice were younger (12 weeks old vs 20 weeks old), and we did not find differences in weight gain, indicating the importance of the age of C57BL/6 mice in our facility in order to develop a stronger metabolic dysfunction phenotype associated with a high-fat diet (11,36).

In parallel, EcN::*ttr* treatment reshaped the bile acid pool, increasing the abundance of SBA Mice fed a high fat diet need to produce more bile acids to support lipid emulsification, which can lead to higher levels of primary bile acids in the large intestine (38). Mice in the EcN::*ttr* group showed lower levels of primary bile acids and precursors, suggesting more efficient ileal reabsorption or better transformation by gut microbiota. Additionally, the EcN*::ttr* group showed reduced levels of diarrhea-associated, sulfated, and glucuronidated SBA, suggesting their gut microbiota may have a greater ability to modify SBA. Given that bile acids act as signaling molecules associated with changes in neuroinflammation levels and expressed behaviour (13) as part of the gut-liver-brain axis, these changes provide a plausible mechanistic link between gut microbial activity and host metabolic and neurobehavioural outcomes. Clinically, both major depressive disorder and anxiety have been related to reductions in the non-12-hydroxylated primary bile acids which was able to be relieved through the use of antidepressants (39). Administration of EcN::*ttr* worked similarly to the antidepressants by reducing the self-soothing behaviour. EcN::*ttr* was also successful in mitigating the long-term memory dysfunction observed, indicating a potential restoration of bile acid function, similar to other studies (40). Bile acid signaling has been implicated in regulating glucose metabolism, inflammation, and central nervous system function, suggesting that EcN::*ttr* may exert pleiotropic effects through coordinated metabolic signaling pathways.

A major limitation in the field of microbiome therapeutics, may be the lack of microbial persistence has hindered clinical translation. By enabling targeted activity within inflamed tissues, engineered live biotherapeutics such as EcN::*ttr* have the potential to deliver more consistent and durable therapeutic effects. Importantly, this approach may extend beyond intestinal diseases to systemic conditions linked to microbiome dysfunction, including metabolic and neurobehavioural disorders. Despite these promising findings, several limitations should be considered. While associations between microbial, metabolic, and behavioural changes are evident, direct causal mechanisms remain to be fully elucidated. Future studies incorporating mechanistic dissection of bile acid signaling pathways, neural circuitry, and immune modulation will be important to define the pathways linking EcN::*ttr* activity to host outcomes. In addition, translation to human populations will require careful evaluation of safety, persistence, and efficacy in the context of diverse diets and disease states.

In conclusion, this study provides evidence that an engineered live biotherapeutic can restore gut–liver–brain axis function under diet-induced stress by leveraging inflammation-associated niches. These findings support a new paradigm in which microbiome-based therapies are designed not only to modulate microbial composition, but to functionally integrate with host physiology in a context-dependent manner. Such approaches may represent a next generation of therapeutics for complex chronic diseases driven by interactions between diet, the microbiome, and host systems.

**Table 1.**
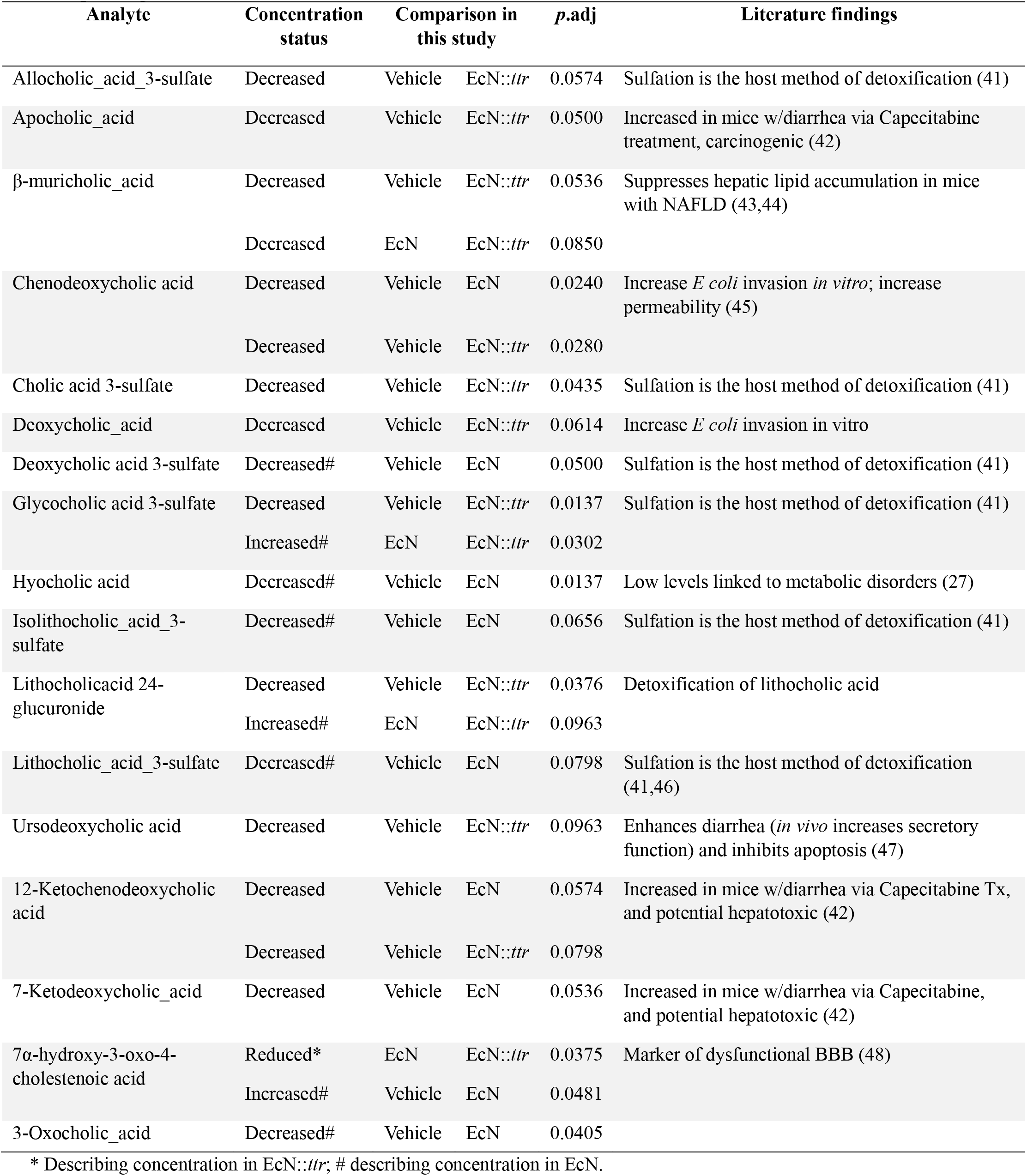
Cecal bile acids modified by EcN::*ttr* administration in mice fed a high omega-6 diet with corresponding actions noted in literature.

| Analyte | Concentration status | Comparison in this study |  | p.adj | Literature findings |
| --- | --- | --- | --- | --- | --- |
| Allocholic_acid_3-sulfate | Decreased | Vehicle | EcN::*ttr* | 0.0574 | Sulfation is the host method of detoxification (41) |
| Apocholic_acid | Decreased | Vehicle | EcN::*ttr* | 0.0500 | Increased in mice w/diarrhea via Capecitabine treatment, carcinogenic (42) |
| β-muricholic_acid | Decreased | Vehicle | EcN::*ttr* | 0.0536 | Suppresses hepatic lipid accumulation in mice with NAFLD (43,44) |
|  | Decreased | EcN | EcN::*ttr* | 0.0850 |  |
| Chenodeoxycholic acid | Decreased | Vehicle | EcN | 0.0240 | Increase *E coli* invasion *in vitro*; increase permeability (45) |
|  | Decreased | Vehicle | EcN::*ttr* | 0.0280 |  |
| Cholic acid 3-sulfate | Decreased | Vehicle | EcN::*ttr* | 0.0435 | Sulfation is the host method of detoxification (41) |
| Deoxycholic_acid | Decreased | Vehicle | EcN::*ttr* | 0.0614 | Increase *E coli* invasion in vitro |
| Deoxycholic acid 3-sulfate | Decreased# | Vehicle | EcN | 0.0500 | Sulfation is the host method of detoxification (41) |
| Glycocholic acid 3-sulfate | Decreased | Vehicle | EcN::*ttr* | 0.0137 | Sulfation is the host method of detoxification (41) |
|  | Increased# | EcN | EcN::*ttr* | 0.0302 |  |
| Hyocholic acid | Decreased# | Vehicle | EcN | 0.0137 | Low levels linked to metabolic disorders (27) |
| Isolithocholic_acid_3-sulfate | Decreased# | Vehicle | EcN | 0.0656 | Sulfation is the host method of detoxification (41) |
| Lithocholicacid 24-glucuronide | Decreased | Vehicle | EcN::*ttr* | 0.0376 | Detoxification of lithocholic acid |
|  | Increased# | EcN | EcN::*ttr* | 0.0963 |  |
| Lithocholic_acid_3-sulfate | Decreased# | Vehicle | EcN | 0.0798 | Sulfation is the host method of detoxification (41,46) |
| Ursodeoxycholic acid | Decreased | Vehicle | EcN::*ttr* | 0.0963 | Enhances diarrhea (*in vivo* increases secretory function) and inhibits apoptosis (47) |
| 12-Ketochenodeoxycholic acid | Decreased | Vehicle | EcN | 0.0574 | Increased in mice w/diarrhea via Capecitabine Tx, and potential hepatotoxic (42) |
|  | Decreased | Vehicle | EcN::*ttr* | 0.0798 |  |
| 7-Ketodeoxycholic_acid | Decreased | Vehicle | EcN | 0.0536 | Increased in mice w/diarrhea via Capecitabine, and potential hepatotoxic (42) |
| 7α-hydroxy-3-oxo-4-cholestenoic acid | Reduced* | EcN | EcN::*ttr* | 0.0375 | Marker of dysfunctional BBB (48) |
|  | Increased# | Vehicle | EcN | 0.0481 |  |
| 3-Oxochoolic_acid | Decreased# | Vehicle | EcN | 0.0405 |  |

\* Describing concentration in EcN::*ttr*; # describing concentration in EcN.

## CRediT Taxonomy

Conceptualization: SG & DLG

Methodology: AV, JKJ, HD, ADH, RI, SG

Visualization: AV, JKJ, EY, DLG

Investigation: AV, JKJ, ADH

Funding acquisition: DLG

Project administration: DLG

Supervision: SG, DLG

Writing – original draft: AV, JKJ

Writing – review & editing: AV, JKJ, SG, DLG

## Data Availability

Data can be found at https://osf.io/h9vdr/overview

## Acknowledgments

We acknowledge U of Vic core metabolomics center for bile acid data generation. We acknowledge Melius MicroBiomics staff Dr. Kevin Horgan (Head of R&D), and other company representatives have approved the manuscript according to the licensing agreement.

## Supplemental Information

**Supplemental Table 1:**
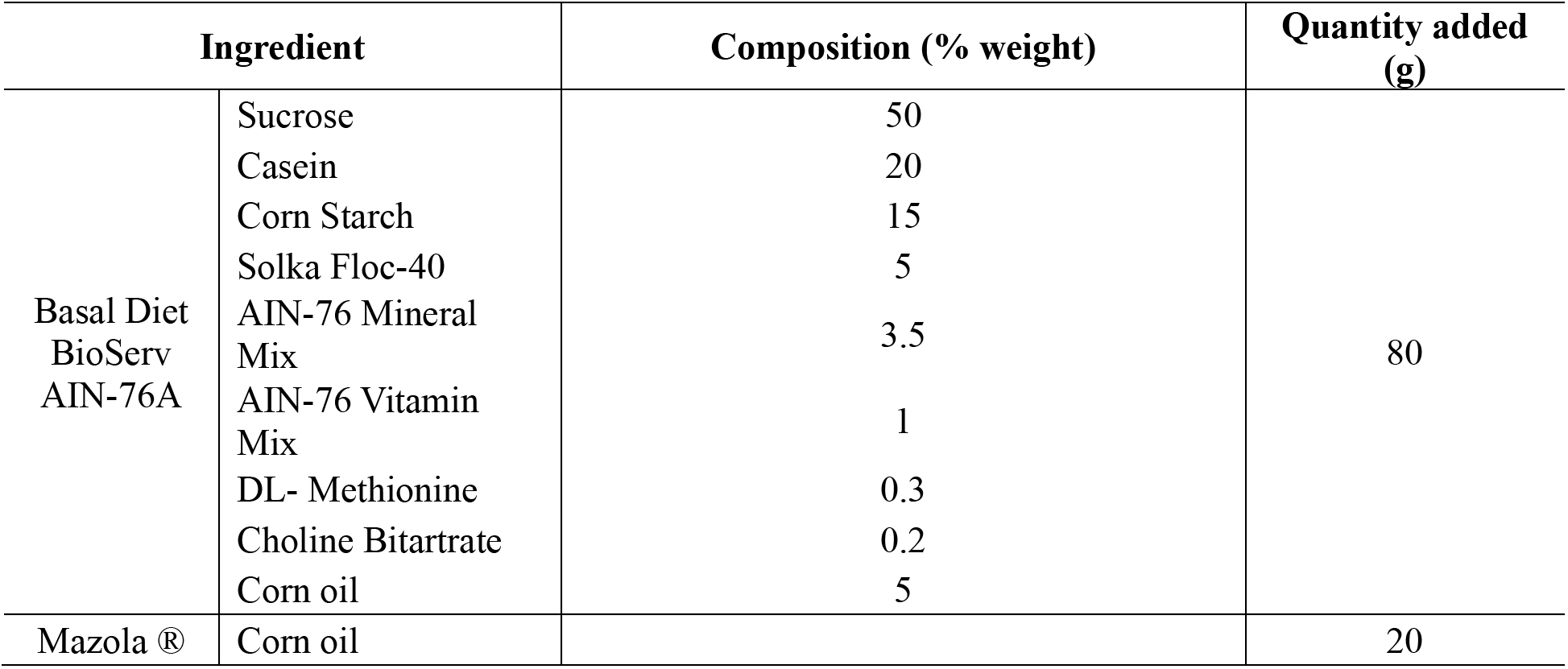
High corn oil diet composition.

**Supplemental Table 2:**
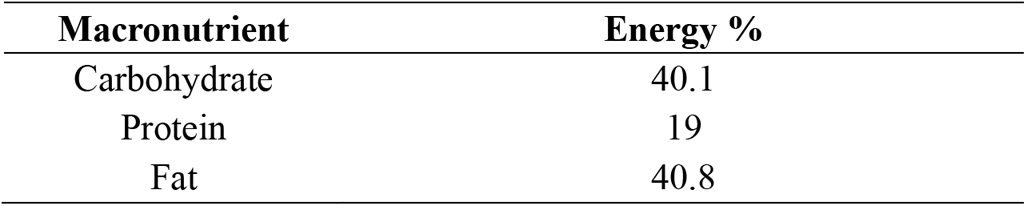
High corn oil diet caloric content.

**Supplemental Table 3:**
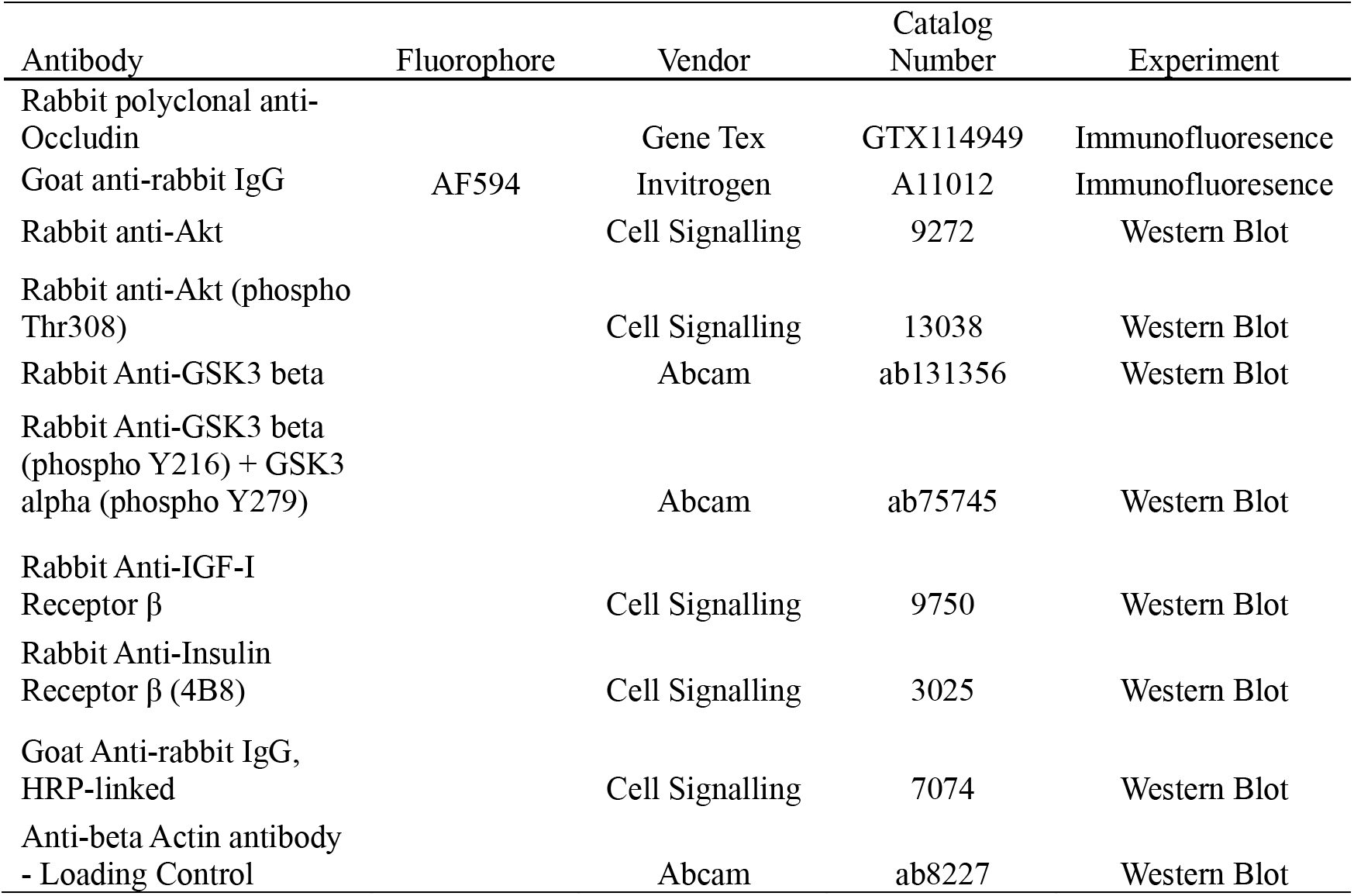
Antibodies used for immunofluorescence and western blots.

**Supplemental Table 4:**
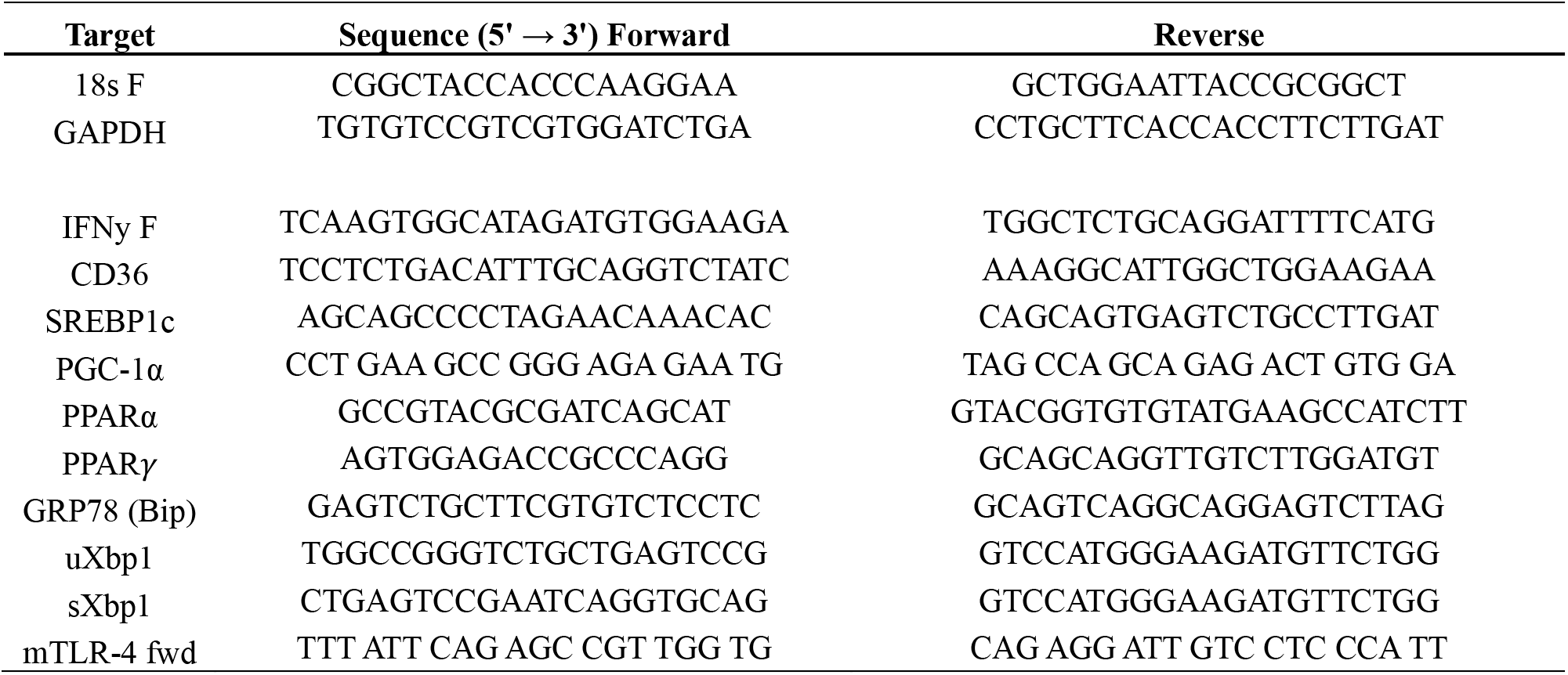
Primers used for qPCR.

**Supplemental Figure 1:**
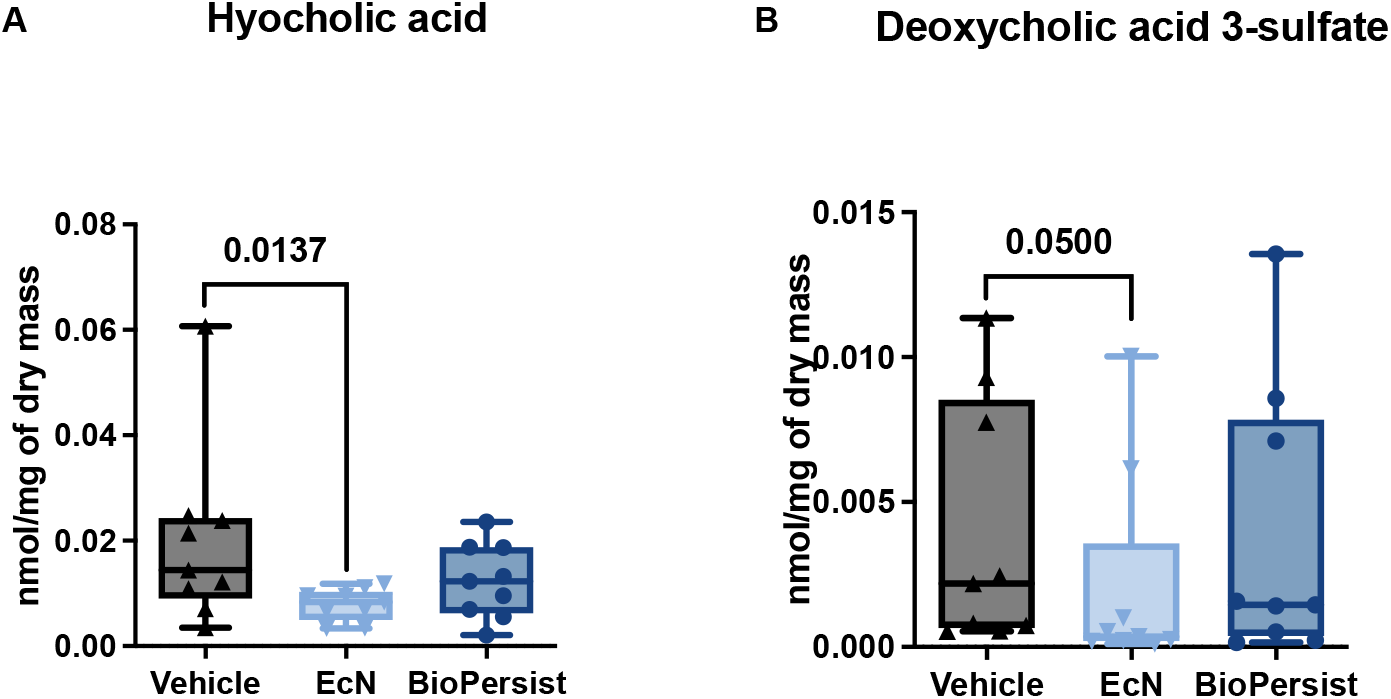
EcN altered bile acid concentrations in mice fed a high omega-6 PUFA diet. Bile acid concentrations (nmol/mg of dry mass) isolated from cecal contents (n=9 per group) with changes highlighted in (A) hyocholic acid and (B) deoxycholic acid 3-sulfate. Data was analysed using Kruskal-Wallis with post-hoc Dunn’s multiple comparisons test. Adjusted *P*-values are shown in the graph.

