## Supplemental Information for "An Engineered Live Biotherapeutic Restores Gut–Liver–Brain Axis Function by Reducing Inflammatory Pathways from Diet-Induced Stress"

#Co-first authors

***Corresponding authors**

**Funding:** SECIHTI for trainee funding (AV), Crohn's and Colitis Canada (DLG), Health Research BC (DLG)

**CRediT Taxonomy:**
Conceptualization: SG & DLG
Methodology: AV, JKJ, HD, ADH, RI, SG
Visualization: AV, JKJ, EY, DLG
Investigation: AV, JKJ, ADH
Funding acquisition: DLG
Project administration: DLG
Supervision: SG, DLG
Writing – original draft: AV, JKJ
Writing – review & editing: AV, JKJ, SG, DLG

**Competing interests:** EcN::*ttr* is patented to The University of British Columbia (US11684643) and licensed to Melius Microbiomics Ltd (MMB; https://www.mmblivebio.com/), a UBC spin-off company co-founded by DLG who served as the Chief Scientific Officer from 10/2022-06/2025 and is currently the non-executive Chief Science Advisor and SG was an advisor from 04/2023-06/2025. SG and DLG have founding shares and options in MMB. AV, JKJ, HD, ADH, RI declare that they have no competing interests.

**Data Availability:** Data can be found at https://osf.io/h9vdr/overview

### Supplemental Table 1: High corn oil diet composition

| Ingredient | | Composition (% weight) | Quantity added (g) |
|---|---|---|---|
| Basal Diet BioServ AIN-76A | Sucrose | 50 | 80 |
| | Casein | 20 | |
| | Corn Starch | 15 | |
| | Solka Floc-40 | 5 | |
| | AIN-76 Mineral Mix | 3.5 | |
| | AIN-76 Vitamin Mix | 1 | |
| | DL- Methionine | 0.3 | |
| | Choline Bitartrate | 0.2 | |
| | Corn oil | 5 | |
| Mazola ® | Corn oil | | 20 |

### Supplemental Table 2: High corn oil diet caloric content

| Macronutrient | Energy % |
|---|---|
| Carbohydrate | 40.1 |
| Protein | 19 |
| Fat | 40.8 |

### Supplemental Table 3: Antibodies used for immunofluorescence and western blots

| Antibody | Fluorophore | Vendor | Catalog Number | Experiment |
|---|---|---|---|---|
| Rabbit polyclonal anti-Occludin | | Gene Tex | GTX114949 | Immunofluoresence |
| Goat anti-rabbit IgG | AF594 | Invitrogen | A11012 | Immunofluoresence |
| Rabbit anti-Akt | | Cell Signalling | 9272 | Western Blot |
| Rabbit anti-Akt (phospho Thr308) | | Cell Signalling | 13038 | Western Blot |
| Rabbit Anti-GSK3 beta | | Abcam | ab131356 | Western Blot |
| Rabbit Anti-GSK3 beta (phospho Y216) + GSK3 alpha (phospho Y279) | | Abcam | ab75745 | Western Blot |
| Rabbit Anti-IGF-I Receptor β | | Cell Signalling | 9750 | Western Blot |
| Rabbit Anti-Insulin Receptor β (4B8) | | Cell Signalling | 3025 | Western Blot |
| Goat Anti-rabbit IgG, HRP-linked | | Cell Signalling | 7074 | Western Blot |
| Anti-beta Actin antibody - Loading Control | | Abcam | ab8227 | Western Blot |

### Supplemental Table 4: Primers used for qPCR

| Target | Sequence (5' → 3') Forward | Reverse |
|---|---|---|
| 18s F | CGGCTACCACCCAAGGAA | GCTGGAATTACCGCGGCT |
| GAPDH | TGTGTCCGTCGTGGATCTGA | CCTGCTTCACCACCTTCTTGAT |
| IFNy F | TCAAGTGGCATAGATGTGGAAGA | TGGCTCTGCAGGATTTTCATG |
| CD36 | TCCTCTGACATTTGCAGGTCTATC | AAAGGCATTGGCTGGAAGAA |
| SREBP1c | AGCAGCCCCTAGAACAAACAC | CAGCAGTGAGTCTGCCTTGAT |
| PGC-1α | CCT GAA GCC GGG AGA GAA TG | TAG CCA GCA GAG ACT GTG GA |
| PPARα | GCCGTACGCGATCAGCAT | GTACGGTGTGTATGAAGCCATCTT |
| PPAR$\gamma$ | AGTGGAGACCGCCCAGG | GCAGCAGGTTGTCTTGGATGT |
| GRP78 (Bip) | GAGTCTGCTTCGTGTCTCCTC | GCAGTCAGGCAGGAGTCTTAG |
| uXbp1 | TGGCCGGGTCTGCTGAGTCCG | GTCCATGGGAAGATGTTCTGG |
| sXbp1 | CTGAGTCCGAATCAGGTGCAG | GTCCATGGGAAGATGTTCTGG |
| mTLR-4 fwd | TTT ATT CAG AGC CGT TGG TG | CAG AGG ATT GTC CTC CCA TT |

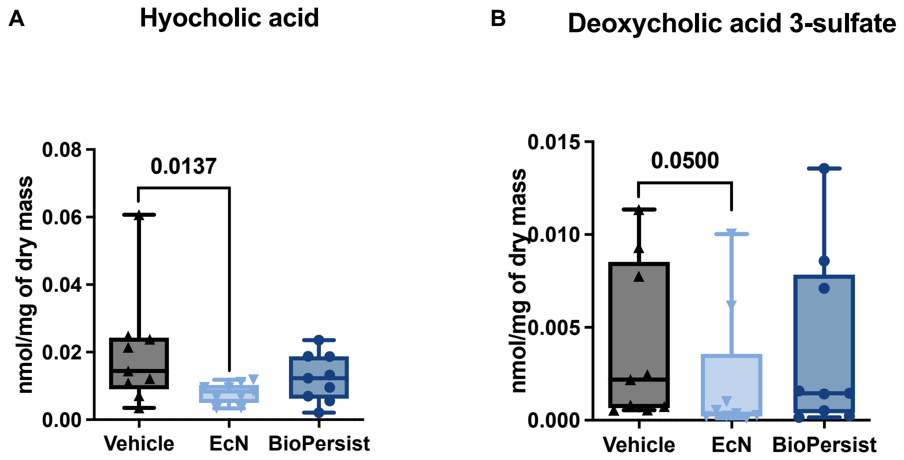


**Supplemental Figure 1: EcN altered bile acid concentrations in mice fed a high omega-6 PUFA diet**

Bile acid concentrations (nmol/mg of dry mass) isolated from cecal contents (n=9 per group) with changes highlighted in (A) hyocholic acid and (B) deoxycholic acid 3-sulfate. Data was analysed using Kruskal-Wallis with post-hoc Dunn's multiple comparisons test. Adjusted *P*-values are shown in the graph.
